# Hijacking SARS-Cov-2/ACE2 receptor interaction by natural and semi-synthetic steroidal agents acting on functional pockets on receptor binding region

**DOI:** 10.1101/2020.06.10.144964

**Authors:** Adriana Carino, Federica Moraca, Bianca Fiorillo, Silvia Marchianò, Valentina Sepe, Michele Biagioli, Claudia Finamore, Silvia Bozza, Daniela Francisci, Eleonora Distrutti, Bruno Catalanotti, Angela Zampella, Stefano Fiorucci

## Abstract

The coronavirus disease 2019 (COVID-19) is a respiratory tract infection caused by the severe acute respiratory syndrome coronavirus (SARS)-CoV-2. In the light of the urgent need to identify novel approaches to be used in the emergency phase, a largely explored strategy has been the repurpose of clinically available drugs as new antivirals, by targeting different viral proteins. In this paper, we describe a drug repurposing strategy based on a virtual screening of druggable pockets located in the central β-sheet core of the SARS-CoV-2 Spike protein RBD supported by in vitro tests identifying several steroidal derivatives as SARS-CoV-2 entry inhibitors. Our results demonstrate that several potential binding sites exist in the SARS CoV-2 S protein, and that the occupancy of these pockets reduces the ability of the S protein RBD to bind to the ACE2 consensus in vitro. In particular, natural occurring and clinically available steroids as glycyrrhetinic and oleanolic acids, as well as the bile acids derivatives glyco-UDCA and obeticholic acid have been shown to be effective in preventing virus entry in the case of low viral load. All together, these results might help to define novel approaches to reduce the viral load by using SARS-CoV-2 entry inhibitors.

## Introduction

The coronavirus disease 2019 (COVID-19) is a respiratory tract infection caused by the severe acute respiratory syndrome (SARS)-CoV-2, a newly emerged coronavirus first identified in the city of Wuhan in China in December 2019 (Zhu et al., 2020). Globally, as of June 9, 2020 there have been more than > 7 million confirmed cases of COVID-19, including 404,396 deaths (World Health Organization, 2020) in 216 countries (Fauci et al., 2020). Genetic sequencing SARS-CoV-2 demonstrates that the virus is a betacoronavirus sharing∼ 80% genetic identity with SARS-CoV and MERS-CoV identified in 2003 and 2012, respectively, and ∼ 96% identity with bat SARS-related CoV (SARS-CoV) RaTG13 (Wang et al., 2020; Wrapp et al., 2020; Yan et al., 2020b). Similarly, to the 2003 and 2012 pandemics caused by these viruses (De Wit et al., 2016), the human infection caused by SARS-CoV-2 induces respiratory symptoms whose severity ranges from asymptomatic/poorly symptomatic to life threatening pneumonia and a cytokine related syndrome that might be fatal (Guan et al., 2020; Zou et al., 2020).

It is well established that similarly to SARS-CoV, the SARS-CoV-2 enters the host cells by hijacking the human angiotensin converting enzyme receptor (ACE2) (Gui et al., 2017; Yuan et al., 2017; Walls et al., 2019, 2020; Wang et al., 2020; Yan et al., 2020a). Interaction of the virus with ACE2 is mediated by the transmembrane spike (S) glycoprotein of SARS-CoV-2, which share e 80% amino acid sequence identity with SARS-Cov and 97.2% sequence homology with the bat SARS-CoV -RaTG13. In the case of SARS-CoV, the spike glycoprotein (S protein) on the virion surface mediates receptor recognition and membrane fusion (Lu et al., 2020). In the intact virus, the S protein assembly in a trimeric structure protruding from the viral surface. Each monomer of the trimeric S protein has a molecular weight of ≈180 kDa and contains two functional subunits, S1 and S2 that mediate, respectively, the attachment to ACE2 and the membrane fusion. The S1 binds to the carboxypeptidase domain of ACE2 with a dissociation constant (Kd) of ∼15 nM (Hoffmann et al., 2020).

Structure analysis has demonstrated that the N- and C-terminal portions of S1 fold as two independent domains, N-terminal domain (NTD) and C-terminal domain (CTD), with this latter corresponding to the receptor-binding domain (RBD) (Wang et al., 2020). According to the high-resolution crystal structure information available thus far, the RBD moves like a hinge between two conformations (‘up’ or ‘down’) to expose or hide the residues that bind the ACE2. Within the RBD, there is a receptor binding motif (RBM) which makes the primary contact with the carboxypeptidase domain of ACE2. Importantly, while amino acid alignment studies have shown that RBD of SARS-CoV-2 shares 73.5% homology with and SARS-CoV, the identity of RBM, the most variable region of RBD, is significantly lower (∼ 50%) making unclear whether the RBMs of the two viruses can induce cross-reactive antibodies. The region outside the RBM is thought to play an important role in maintaining the structural stability of the RBD.

The entry of SARS-CoV-2 in host cells requires the cleavage of the S protein. The cleavage process takes place in two steps. After binding to ACE2, the S protein is cleaved by cellular proteases between the S1 and S2 subunits (Li et al., 2003; Lan et al., 2020; Shang et al., 2020; Wang et al., 2020). This first step is operated by a camostat-sensitive transmembrane serine protease TMPRSS2. The SARS-CoV-2 has a distinct furin cleavage site (Arg-Arg-Ala-Arg) at residues 682–685, between the S1 and S2 domains not found in SARS-CoV, which may explain some of the biological differences. This furin cleavage site expands the versatility of SARS-CoV-2 for cleavage by cellular proteases and potentially increases the tropism and transmissibility owing to the wide cellular expression of furin proteases especially in the respiratory tract (Belouzard et al., 2009; Ou et al., 2020). Cleavage at the S1/S2 site is essential to unlock the S2 subunit which, in turn, drives the membrane fusion. Importantly, a second S2 site of cleavage has been identified at the S2′ site which is thought essential to activate the protein for membrane fusion.

The spreading of the COVID-19 pandemic and the lack of effective therapies targeting the viral replication has prompted an impressive amount of investigations aimed at targeting several aspects of SARS-CoV-2 biology and viral interactions with ACE2. In this scenario, drug repurposing is a well-established strategy to quickly move already approved or shelved drugs to novel therapeutic targets bypassing the time-consuming stages of drug development. This accelerated drug development and validation strategy has led to numerous clinical trials for the treatment of COVID-19 (Li and De Clercq, 2020; Liu et al., 2020). Despite several encouraging results, treatment of SARS-CoV-2 infection remains suboptimal and there is an urgent need to identify novel approaches to be used in the emergency phase.

One of such approaches is to prevent the S protein/ACE2 interaction as a strategy to prevent SARS-CoV-2 entry into target cells. Indeed, and virtual screening campaigns have already identified small molecules able to bind residues at the interface between the RBD of SARS-CoV-2 S protein and the ACE2 receptor (Ghosh et al., 2020; Micholas and Jeremy C., 2020; Senathilake et al., 2020; Utomo et al., 2020; Yan et al., 2020b; Zhou et al., 2020).

In this paper, we have expanded in this area. Our results will demonstrate that, not only several potential binding sites exist in the SARS CoV-2 S protein, but that occupancy of these pockets reduces the ability of the S protein RBD to bind to the ACE2 consensus *in vitro*. Together, these results might help to define novel treatments to reduce the viral load by using SARS-CoV-2 entry inhibitors.

## Material and Methods

### Virtual screening

The electron microscopy (EM) model of SARS-CoV-2 Spike glycoprotein was downloaded from the Protein Data Bank (PDB ID: 6VSB). Missing loops were added by the Swiss-Model web-site (Wrapp et al., 2020). The obtained model was submitted to the Protein Preparation Wizard tool implemented into Maestro ver. 2019 (Schröedinger, 2019) to assign bond orders, adding all hydrogen atoms and adjusting disulfide bonds. The pocket search was performed with the Fpocket website (Schmidtke et al., 2010).

The AutoDock4.2.6 suite (Morris et al., 2009) and the Raccoon2 graphical interface (Forli et al., 2016) were used to carry out the virtual screening approach using the Lamarckian genetic algorithm (LGA). This hybrid algorithm combines two conformational research methods: the genetic algorithm and the local research. For the first low-accuracy screening, for each of the 2906 drugs, 3 poses were generated using 250000 steps of genetic algorithm and 300 steps of local search, while in the second high-accuracy screening protocol, we have generated 20 poses for each ligand, increasing the number of genetic algorithm steps to 25000000. The MGLTools were used to convert both ligands and each pocket into appropriate pdbqt files.

Each of the six selected RBD pocket were submitted to the AutoGrid4 tool, which calculates, for each bonding pocket, maps (or grids) of interaction, considering the different ligands and receptor atom types through the definition of a cubic box. Subsequently, for each grid AutoDock4 calculates interaction energies (ADscore) that express the affinity of a given ligand for the receptor. The library of FDA approved drugs has been obtained both from DrugBank (2106 compounds) (Drugbank, 2020) and from the Selleckchem website (FDA-approved Drug Library, 2020) (tot. 2638). Each database was converted to 3D and prepared with the LigPrep tool (Schröedinger, 2019) considering a protonation state at a physiological pH of 7.4. Subsequently, the two libraries were merged and deduplicated with Open Babel (O’Boyle et al., 2011), giving a total amount of 2906 drugs. The bile acids (BA) focused library was prepared with the same protocol described above. All the images are rendered using UCSF Chimera (Pettersen et al., 2004).

### Chemistry

OCA, BAR704, BAR501 and BAR502 were synthesized as previously described (Festa et al., 2014; Sepe et al., 2016)

### ACE2/SARS-CoV-2 Spike Inhibitor Screening Assay Kit

We tested the selected compounds (UDCA, T-UDCA, G-UDCA, CDCA, G-CDCA, OCA, BAR501, BAR502, BAR704, betulinic acid, oleanolic acid, glycyrrhetinic acid, canrenone Potassium) using the ACE2:SARS-CoV-2 Spike Inhibitor Screening Assay Kit (BPS Bioscience Cat. number #79936) according to the manufacturer’s instructions. All compounds were tested at different concentrations in a range from 0.01 µM to 100 µM. In addition, a concentration-response curve for the Spike protein (0.1-100 nM) was constructed to confirm a concentration-dependent increase in luminescence. A spike concentration of 5 nM was used for the screening of the compounds. Briefly, thaw ACE2 protein on ice and dilute to 1 µg/ml in PBS. Use 50 µL of ACE Solution to coat a 96-well nickel-coated plate and incubate one hour at room temperature with slow shaking. Wash the plate 3 times and incubated 10 minutes with a Blocking Buffer. Next, add 10 µl of inhibitor solution containing the selected compounds and incubated one hour at room temperature with slow shaking. For the “Positive Control” and “Blank”, add 10 µl of inhibitor buffer (5% DMSO solution). After the incubation thaw SARS-CoV-2 Spike (RBD)-Fc on ice and dilute to 0,25 ng/µl (approximately 5 nM) in Assay Buffer 1. Add the diluted Spike protein to each well, except to the Blank. Incubate the reaction 1 hour at room temperature with slow shaking. After 3 washes and incubation with a Blocking Buffer (10 minutes), treat the plate with an Anti-mouse-Fc-HRP and incubated 1 hour at room temperature with slow shaking. Finally, add an HRP substrate to the plate to produce chemiluminescence, which then can be measured using FluoStar Omega microplate reader.

### Quantitative analysis of the anti-SARS-CoV-2 IgG antibodies

Remnant serum samples from post COVID-19 subjects were used. The plasma was collected at the blood bank of Azienda Ospedaliera di Perugia from post COVID-19 donors who participate to “Tsunami” (i.e. TranSfUsion of coNvaleScent plAsma for the treatment of severe pneuMonIa due to SARS.CoV2) program. As per protocol, 1 ml of serum is used for analysis of immunoglobulin concentrations and anti-HCV, anti-HIV, HbsAg and HBsAb. Remnants (aprox. 50 µl) were used in this study. subjects were plasma donors and we The quantitative analysis of the anti-SARS-CoV-2 IgG antibodies directed against the subunits (S1) and (S2) of the virus spike protein were measured by chemiluminescence immunoassay (CLIA) technology (LIAISON®SARS-CoV-2 IgG kit, DiaSorin®, Saluggia, Italy) in serum from COVID-19 PCR-confirmed cases (n = 5) who have recovered from coronavirus disease 2019 (COVID-19) in Italy.

Blood samples were collected in serum collection tubes (BD Vacutainer SST II advance, BD, Plymouth, UK) according to standardized operating procedure. Samples were then centrifuged at 3500 rpm (2451 g) for 10 min. Serum was then collected, and samples were analyzed as soon as possible. The analysis was carried out using the LIAISON®SARS-CoV-2 IgG kit on a LIAISON®XL analyzer in accordance with the manufacturer’s instructions.

## Results

### Virtual Screening of the FDA-approved drug library

With the aim to identify chemical scaffolds capable of inhibiting ACE2/Spike interaction by targeting the RBD of S1 domain of the SARS-CoV-2, we have carried out a virtual screening campaign on an FDA-approved drug library, using the RBD 3D structure obtained from the Protein Data Bank (PDB ID 6SVB; Chain A, residues N331-A520)(Wrapp et al., 2020). Missing regions in the structure were built through the SwissModel webserver (Bertoni et al., 2017). A pocket search with the Fpocket web-server (Le Guilloux et al., 2009) was performed and resulted in the identification of ≈ 300 putative pockets on the whole trimeric structure of the S protein. This search was further refined to identify selected pockets only in the RBD according to three main factors: i) the potential druggability, i.e. the possibility of interfering, directly or through an allosteric mechanism, with the interaction with ACE2; ii) the flexibility degree of the pockets, i.e. excluding pockets defined, even partially, by highly flexible loops whose coordinates were not defined in the experimental structure; iii) sequence conservation with respect to SARS-CoV RBD (Figure 1A). On these bases, 6 pockets have been selected on the RBD and numbered according to Fpocket ranking (Figure 1 A and C).

**Figure 1.**
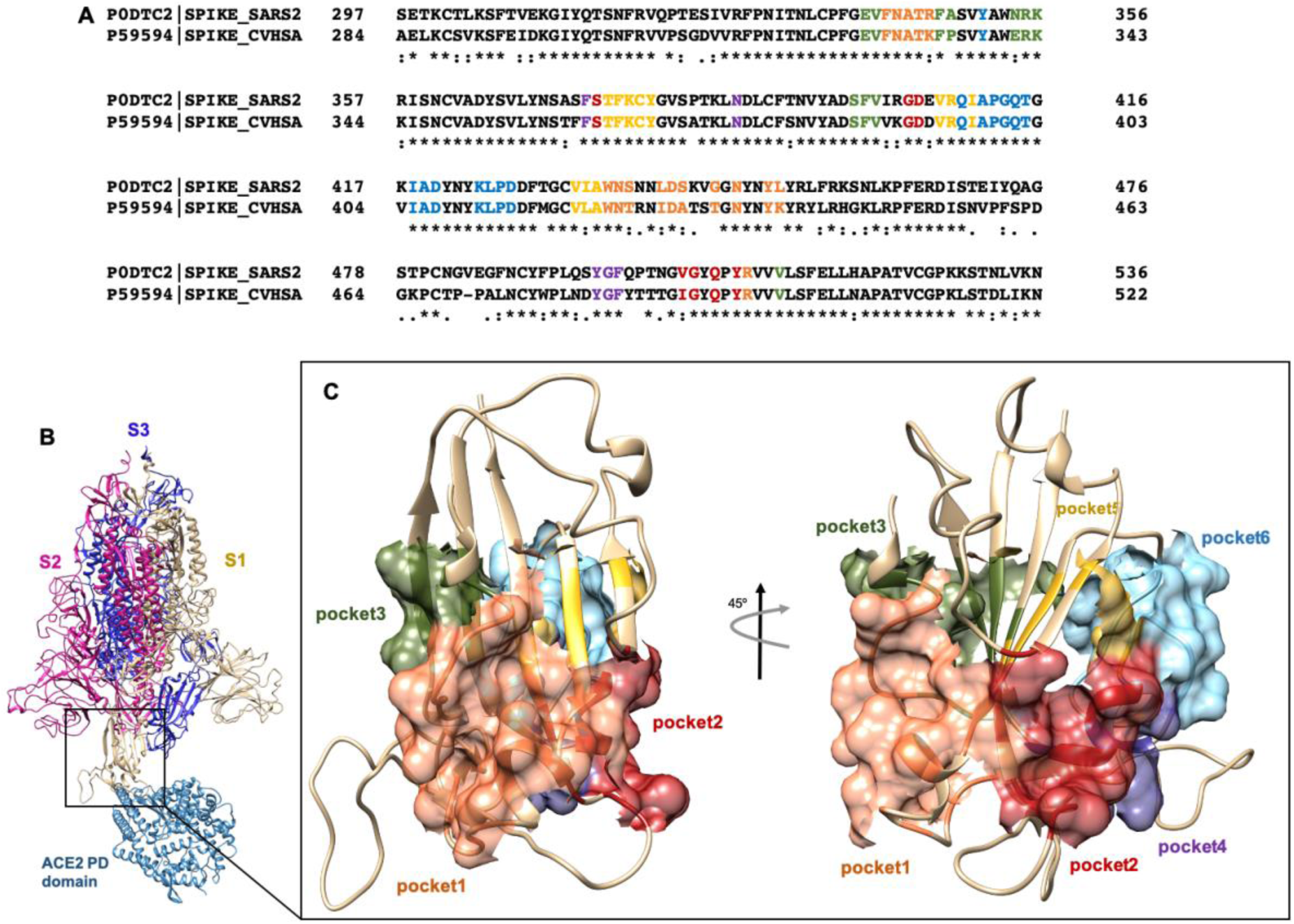
(A) Clustal Omega alignment of RBD regions of SARS-CoV and SARS-CoV-2 Spike protein. Residues bearing to different pockets are colored respectively yellow (Pocket 1), green (Pocket 2), light blue (Pocket 3), magenta (Pocket 4), red (pocket 5) and dark slate blue (Pocket 6). (B) Cartoon representation of the trimer of SARS-2 Spike protein in complex with the PD domain of ACE2. Complex obtained through the superposition of the PDB structures 6VSB and 6M0J. (C) Surface representation of the six selected pockets used for the screening.

**Figure 2.**
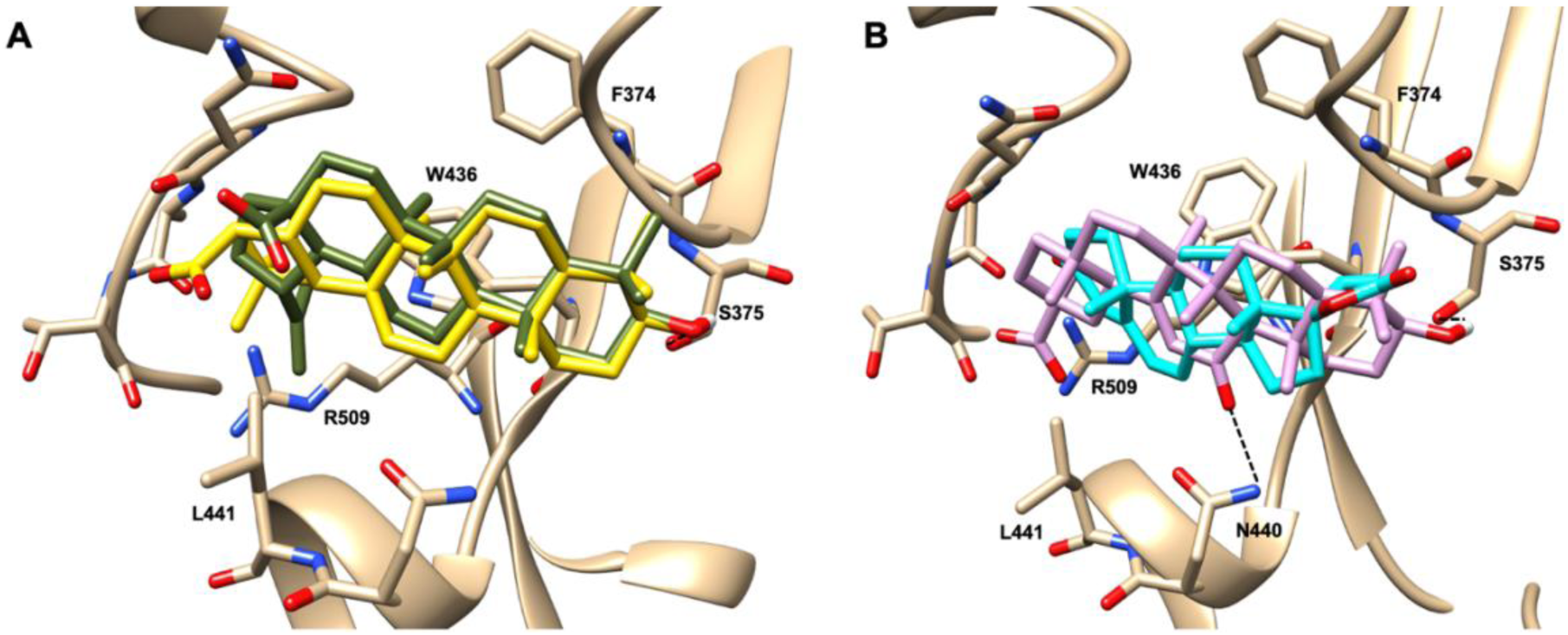
Graphical representation of the binding mode of the best compounds resulting from the screening in pocket 1. The RBD region is represented in transparent surface colored by residues hydrophobicity. Color codes are: dodger blue for the most hydrophilic regions, white, to orange-red for the most hydrophobic. (**A**) Betulinic acid (dark olive-green stick) and oleanolic acid (gold stick). (**B**) Glycyrrhetinic acid (plum stick) and potassium canrenoate (cyan stick). For clarity reasons hydrogen atoms are omitted and only interacting aminoacids are displayed in sticks.

First, these pockets were used for the virtual screening of 2906 FDA-approved drugs from the DrugBank and the Selleckchem websites, using the AutoDock4.2.6 program (Morris et al., 2009) and the Raccoon2 graphical user interface (Forli et al., 2016). This step was followed by a high-accuracy, screening based on the binding affinity predicted by AutoDock4 (ADscore), focusing our investigation on the results showing an ADscore lower than -6 kcal/mol.

These studies allowed the identification of several compounds with steroidal and triterpenoidal scaffold, including glycyrrhetinic acid, betulinic acid and the corresponding alcohol (betulin), canrenone and the corresponding opened form on the γ-lactone ring as potassium salt (potassium canrenoate), spironolactone and oleanolic acid, showing robust binding selectivity towards the RBD’s pocket 1 (Table 1).

**Table 1.**
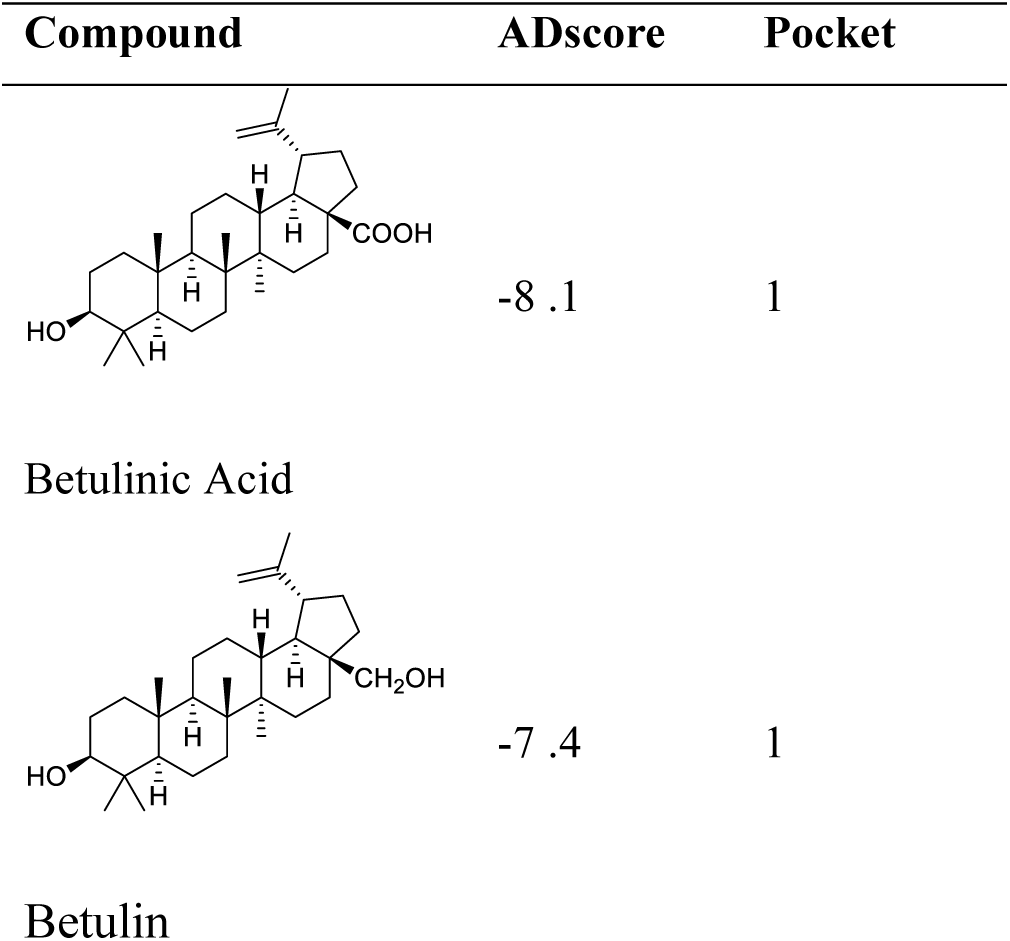
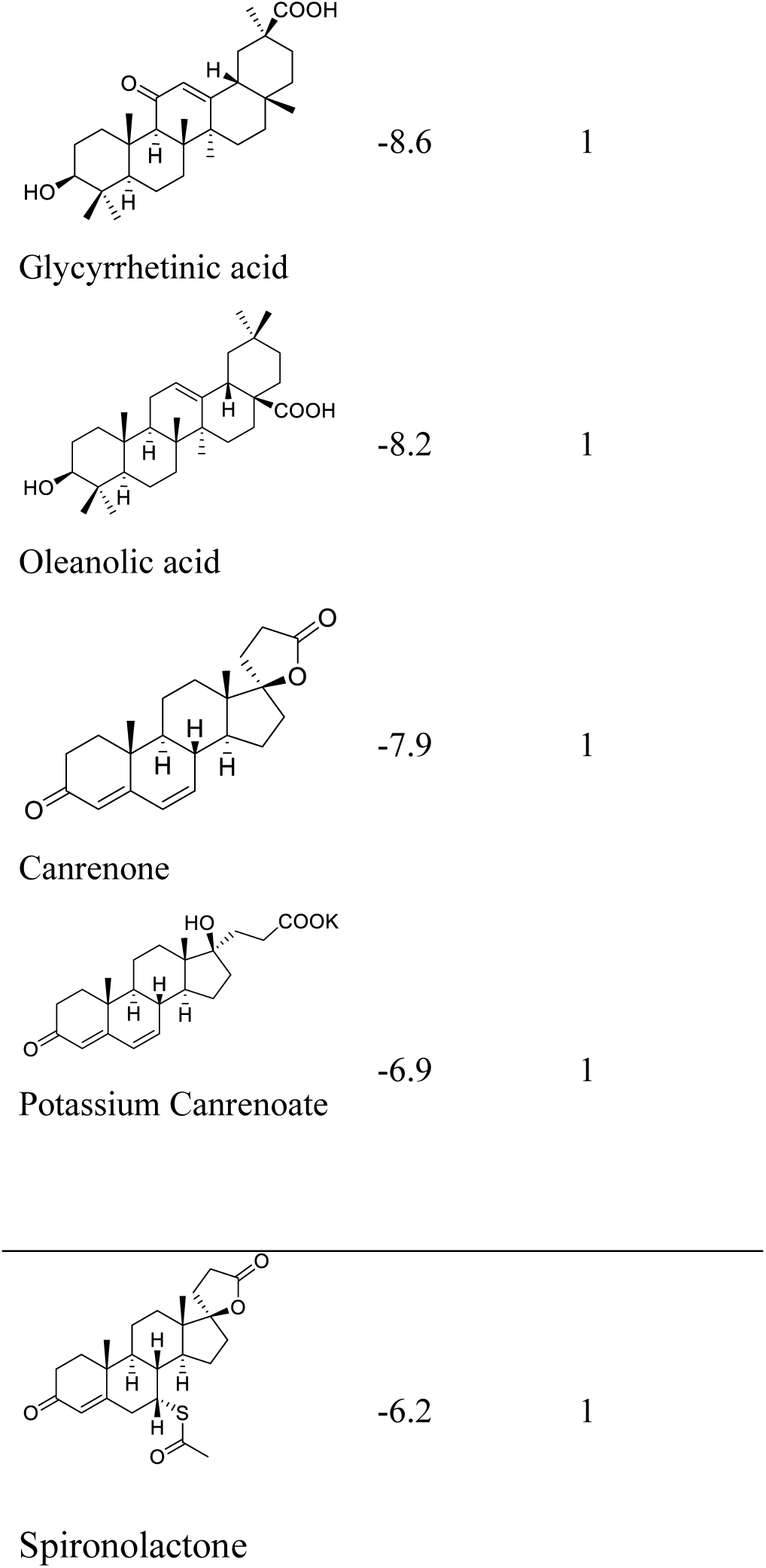
Results of the screening of FDA approved drugs on the RBD region of the Spike protein of SARS-CoV-2 with the Autodock 4.2.6 program. Binding affinity values (ADscore) are expressed in kcal/mol.

RBD’s pocket 1, located on the β-sheet in the central core of the RBD, is the less conserved among the screened, presenting five conservative (R346K, S438T, L440I, S442A) and two non-conservative (G445T and L451K) mutations from SARS-CoV-2 to SARS-CoV.

Glycyrrhetinic acid, the best compound according to the AD score, binds the pocket through both hydrophobic and polar interactions. The steroidal scaffold relied between the hydrophobic side of the β-sheet core of RBD, defined by W436, F374 and the side chain of R509, and L441 on the other side, engaging hydrophobic contacts. In addition, the binding is reinforced by ionic contacts between the carboxyl group with R509, and by hydrogen bonds between the carbonyl group with N440 and the hydroxyl group with S375. Oleanolic acid and betulinic acid showed similar binding modes with the main difference in the carboxylic groups oriented towards the solvent. Finally, potassium canrenoate showed a different orientation of the steroidal system within the binding site, with the carboxylic function weakly bonded to S375 (3.1 Å), and the π-system of rings A and B stacked between W436 and L441.

Because the above mentioned triterpenoids have been identified as natural ligands for two bile acid activated receptors, the Farnesoid-X-Receptor (FXR) and G protein Bile Acid Receptor (GPBAR)-1 (Sepe et al., 2015; De Marino et al., 2019; Fiorucci and Distrutti, 2019), we have further investigated whether mammalian ligands of these receptors were also endowed with the ability to bind to the above mentioned RBD’s pockets. More specifically, oleanolic, betulinic and ursolic acid have been proved to act as selective and potent GPBAR1 agonists (Sato et al., 2007; Genet et al., 2010; Lo et al., 2016), while glycyrrhetinic acid, the major metabolic component of licorice, and its corresponding saponin, glycyrrhizic acid, have been shown to act as dual FXR and GPBAR1 agonists in transactivation assay (Distrutti et al., 2015) promoting GLP-1 secretion in type 1-like diabetic rats (Wang et al., 2017). Bile acids (BA) are steroidal molecules generated in the liver from cholesterol breakdown (Fiorucci and Distrutti, 2019). Primary BA include two main bile acid species, cholic acid (CA) and chenodeoxycholic acid (CDCA), which have been recognized to function as the main FXR ligand in humans (Fiorucci and Distrutti, 2019). Secondary BA, deoxycholic acid and lithocholic acid (DCA and LCA) generated by intestinal microbiota, are preferential ligands for GPBAR1 (Maruyama et al., 2002; Fiorucci and Distrutti, 2019). Ursodeoxycholic acid (UDCA) that is a primary bile acid in mice, but a “tertiary” bile acid found in trace in humans, is along with CDCA the only bile acid approved for clinical use, and is a weak agonist for GPBAR1 and considered a neutral or weak antagonist toward FXR (Carino et al., 2019).

Taking into account the structural similarity and the ability to bind the same receptor systems, we have carried out and in-deep docking analysis of natural BA and their semisynthetic derivatives currently available in therapy or under pre-clinical and clinical development (De Marino et al., 2019) and tested them for their ability to bind on the above-mentioned pockets in the RBD’s of SARS-CoV-2 S protein (Table 2).

**Table 2.**
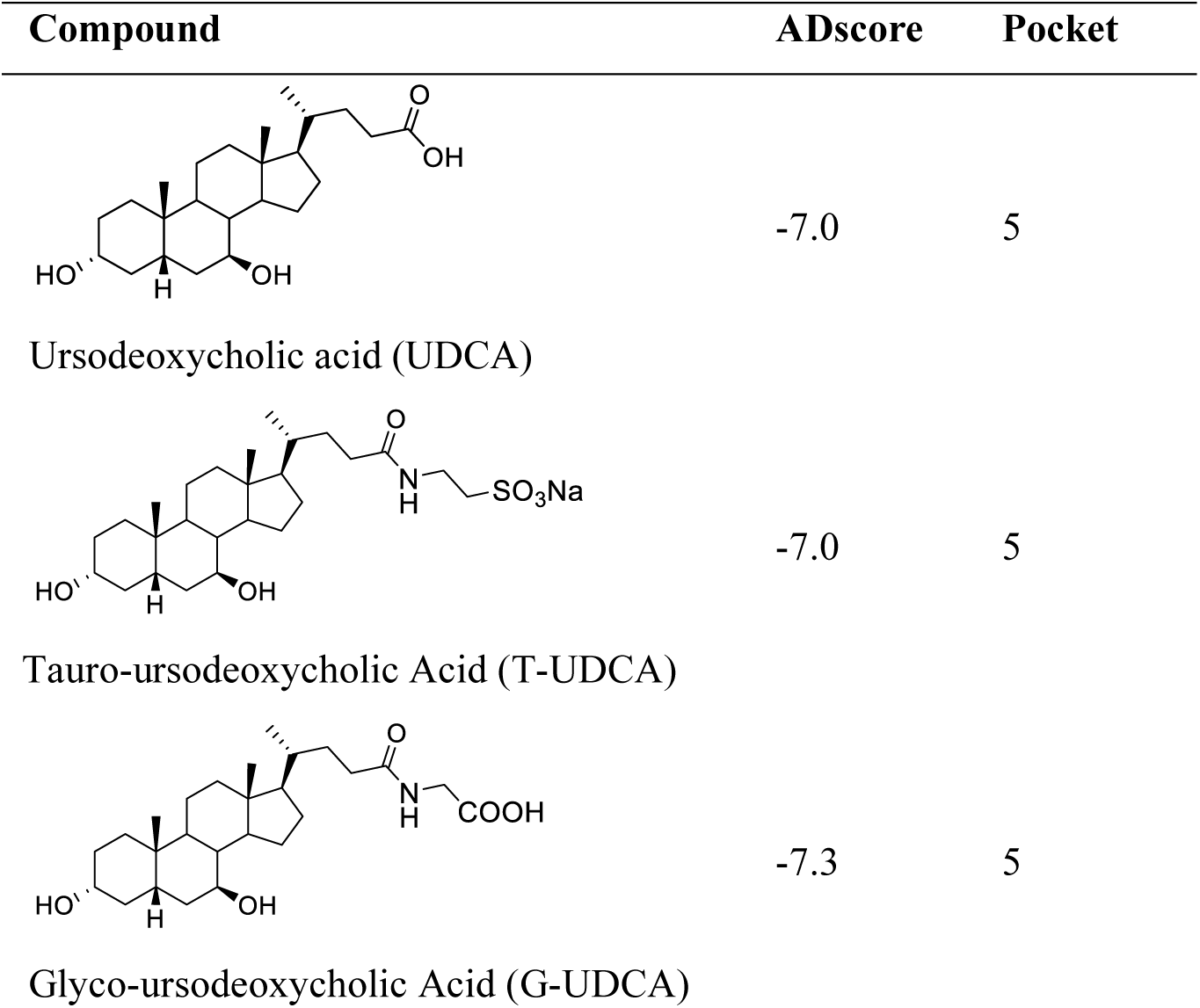
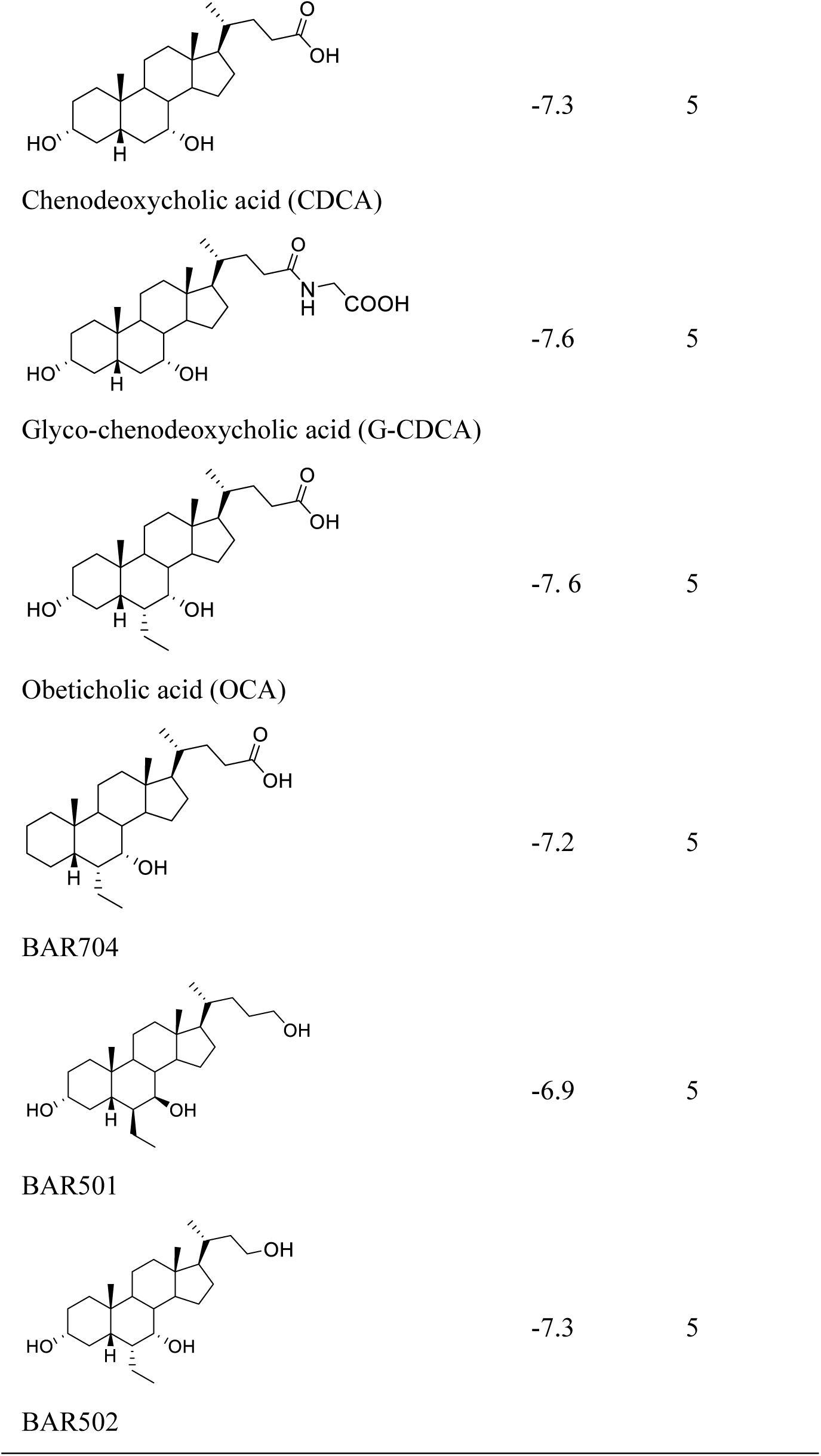
Results of the screening of natural bile acids on the RBD region of the Spike protein of SARS-CoV-2 with the Autodock 4.2.6 program. Binding affinity values (ADscore) are expressed in kcal/mol.

As shown in Table 2, natural BA and their semi-synthetic derivatives exhibit higher affinity scores for the pocket 5. This pocket (Figure 3 A, B and C) included residues bearing to the central β-sheet core but on a different side respect to the pocket 1. The pocket resulted very conserved, showing only one mutation, I434L, from SARS-CoV-2 to SARS-CoV.

**Figure 3.**
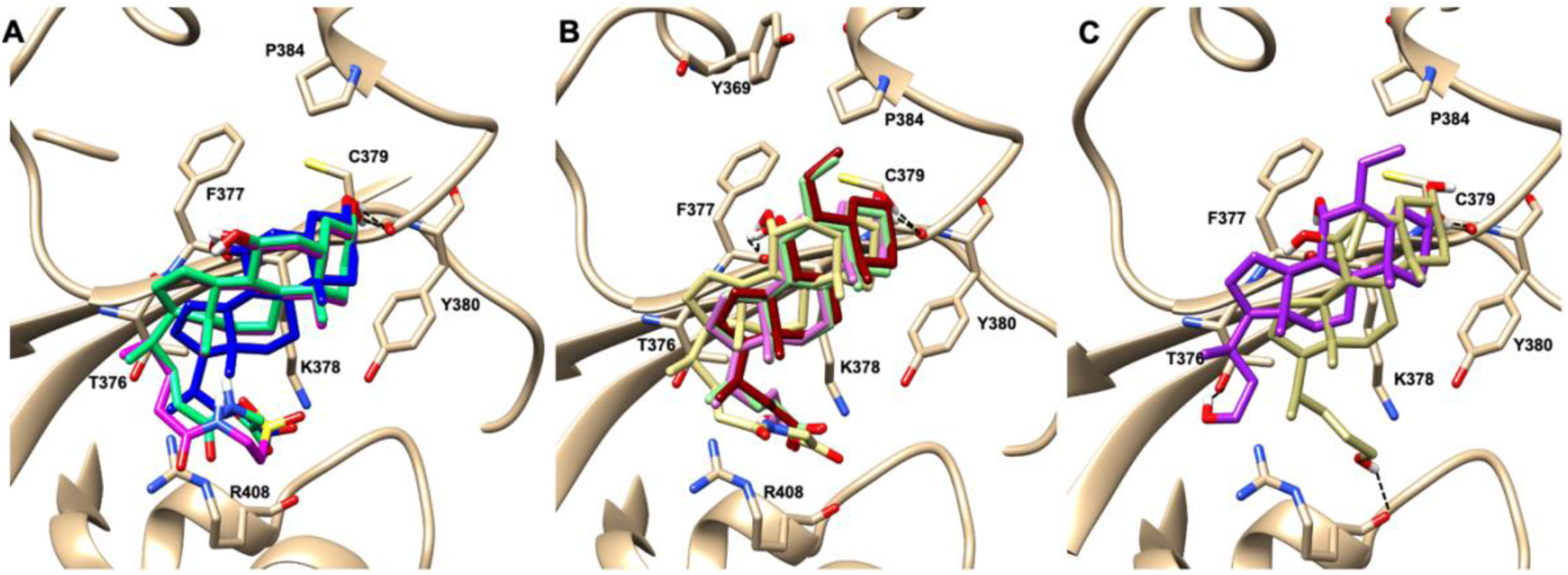
Graphical representation of the binding mode of the best compounds resulting from the screening in pocket 5. The RBD region is represented in tan cartoon, while the pocket 5 residues as transparent surface colored by residues hydrophobicity. Color codes are: dodger blue for the most hydrophilic regions, white, to orange-red for the most hydrophobic. (**A**) UDCA (blue stick), T-UDCA (magenta stick) and G-UDCA (spring-green stick); (**B**) CDCA (orchid stick), OCA (light-green stick), BAR704 (dark-red stick) and G-CDCA (khaki stick); (**C**) BAR501 (gold stick) and BAR502 (purple stick). For clarity reasons hydrogen atoms are omitted and only interacting aminoacids are displayed in sticks.

In the binding mode of UDCA, the carboxylic group on the side chain is positioned between K378 and R408 and the steroidal scaffold is placed in a hydrophobic surface defined by the side chain of K378, T376, F377, Y380 and P384. Additionally, the 3 β-hydroxyl group on ring A H-bonds with the backbone carbonyl of C379. The corresponding glycine and taurine-conjugated derivatives (G-UDCA and T-UDCA, respectively) showed the same ionic interactions of their negatively charged groups with K378 and R408, and the H-bond with the backbone carbonyl of C379 albeit the greater length of the side chain induces a shift of the steroidal system to T376, and an additional π-interaction between the electron density of the glycine amide region and the guanidine moiety of R408, resulting in a better score for G-UDCA, and a reduction in the case of T-UDCA, likely due to a non-optimal arrangement of the taurine moiety within the binding pocket. CDCA showed a very similar binding mode with the only difference in gaining an additional H-bond with the backbone carbonyl of F377 due to the modification in the configuration of the C-7 hydroxyl group (α-oriented in CDCA and β-oriented in UDCA). As for G-UDCA, also G-CDCA established the same H-bonds network of the parent CDCA, while the steroidal core slightly shifted as described for G-UDCA. Interestingly, ADscores of G-UDCA and G-CDCA clearly indicated that the H-bond between the hydroxyl group at C-7 and F377 doesn’t contribute significantly to the binding mode.

Respect to CDCA, the introduction of the ethyl group at the C-6 position as in OCA and in BAR704 improves the internal energy of the ligand (−0.27 for CDCA *vs* -0.59 and -0.60 kcal/mol for OCA and BAR704, respectively) and further favors the binding (Figure 3B), even if the 6-ethyl group, albeit in close proximity of P384 and Y369, did not show any particular contact within the RBD region. BAR501, a neutral UDCA derivative, with an alcoholic side chain end group and the 6 β-ethyl group showed a very similar binding mode compared to the parent compound, with the side chain hydroxyl group H-bonded to R408. Finally, BAR502, with a one carbon less on the side chain positioned the steroidal core as for G-CDCA, thus allowing the C-23 OH group H-bonding with the side chain hydroxyl group of T376.

### *In vitro* screening

Given the results of the virtual screening, we have then investigated whether the agents mentioned in Table 1 and 2 impact on the binding of S protein to the ACE2 receptor. For this purpose, a Spike/ACE2 Inhibitor Screening Assay Kit was used. The assay is designed for screening and profiling inhibitors RBD/ACE2 interaction. To validate the assay, we have first performed a concentration-response curve by adding increasing concentrations of the Spike RBD (0.1-100 nM) and confirmed a concentration-dependent increase of luminescence (n=5 experiments, Figure 4A). Since the curve was linear in the range from 0.1 to 10 nM we have used the concentration of 5 nM for all the following assays. As illustrated in Figure 4, we have found that incubating the Spike RBD with betulinic acid, glycyrrhetinic acid, oleanolic acid and potassium canrenoate (the active metabolite of spironolactone) results in concentration dependent reductions of the binding of S Spike RBD to the ACE2 receptors. While all agents effectively reversed the binding at a concentration of 10 µM, betulinic acid and oleanolic acid showed a significant inhibition at a concentration of 0.1 and 1 µM, respectively (n=of al at least experiments).

**Figure 4.**
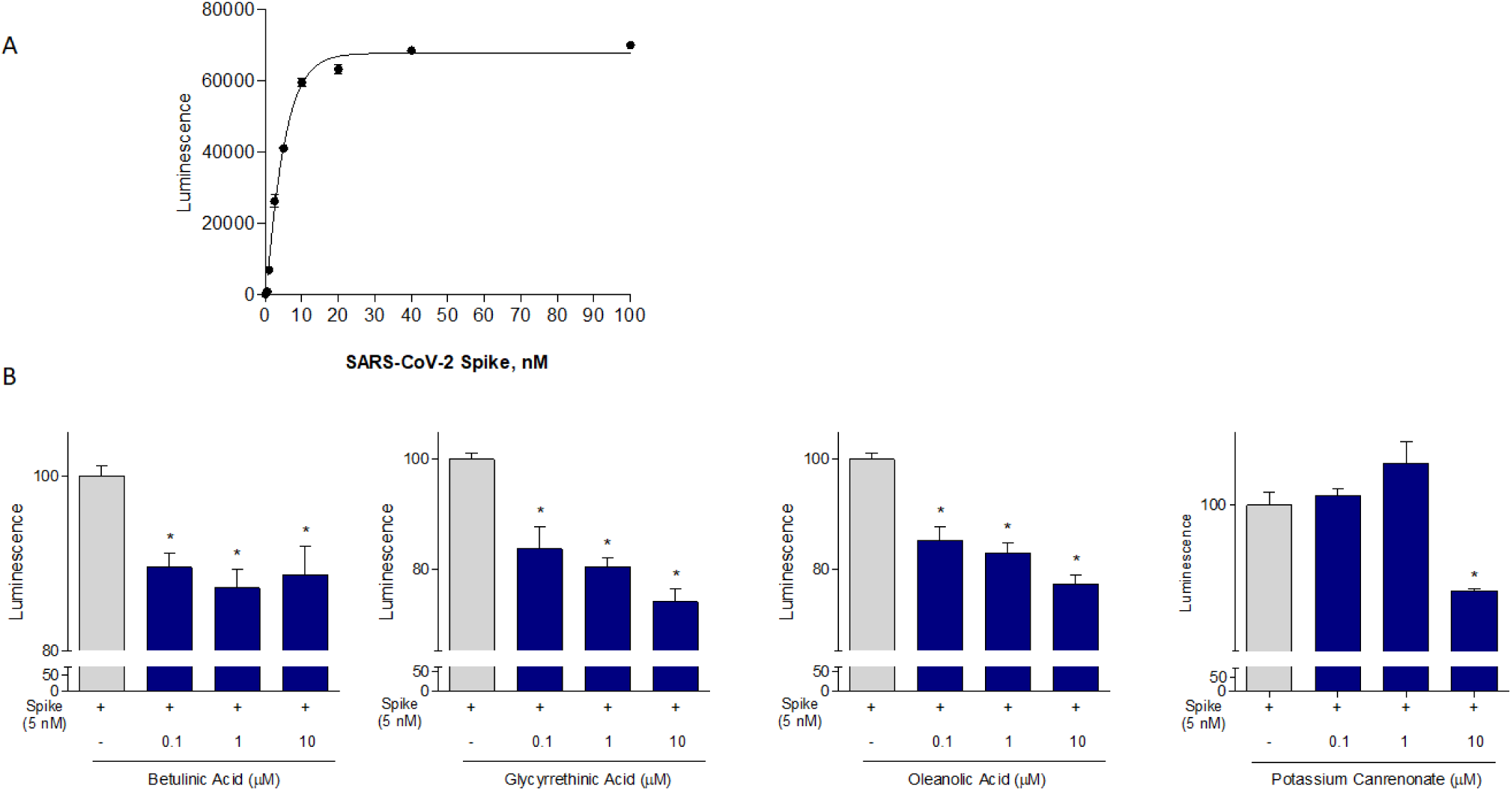
The ACE2:SARS-CoV-2 Spike Inhibitor Screening assay was performed as described in Material and Method section. Data shown are: (**A**) SARS-Cov-2 Spike binding to immobilized ACE2, using an increasing dose of Spike protein (0,5-100 nM); Luminescence was measured using a Fluo-Star Omega fluorescent microplate reader. (**B**) Betulinc acid, glycyrrethinic acid, oleanolic acid and potassium canrenonate were tested at different concentration (0.1, 1 and 10 µM), to evaluate their ability to inhibit the binding of Spike protein (5 nM) to immobilized ACE2, by using the ACE2:SARS-CoV-2 Spike Inhibitor Screening assay Kit. Luminescence was measured using a Fluo-Star Omega fluorescent microplate reader. Luminescence values of Spike 5 nM were arbitrarily set to 100 %. Results are expressed as mean ± standard error. *p < 0.05 versus Spike 5 nM. Data are the mean ± SE, n = 3.

Because these data demonstrate that betulinic acid and oleanolic acid were effective in inhibiting the bind of S protein’s RBD to ACE2, and the two triterpenoids were known for their ability to bind to GPBAR1, a bile acid activated receptor, we then tested whether natural bile acids were also effective in reducing the SARS-CoV-2-ACE2 interaction. As illustrated in Figure 5, the secondary bile acid UDCA and its taurine conjugate, T-UDCA, caused a slight and dose dependent inhibition of S protein RBD to the ACE2 receptor (Figure 5 A and B). In contrast, G-UDCA, i.e. the main metabolite of UDCA in human, inhibits the RBD binding to the ACE2 receptor by approx. 50% in a concentration dependent manner. Similar concentration dependent effects were observed with CDCA (Figure 5D). The CDCA metabolite, G-CDCA exerted a similar inhibitory effect with 0.1 µM being the most effective concentration. A combination of UDCA and G-CDCA gives the same amount of inhibition than G-CDCA alone confirming that UDCA itself as a very limited inhibitory activity.

**Figure 5.**
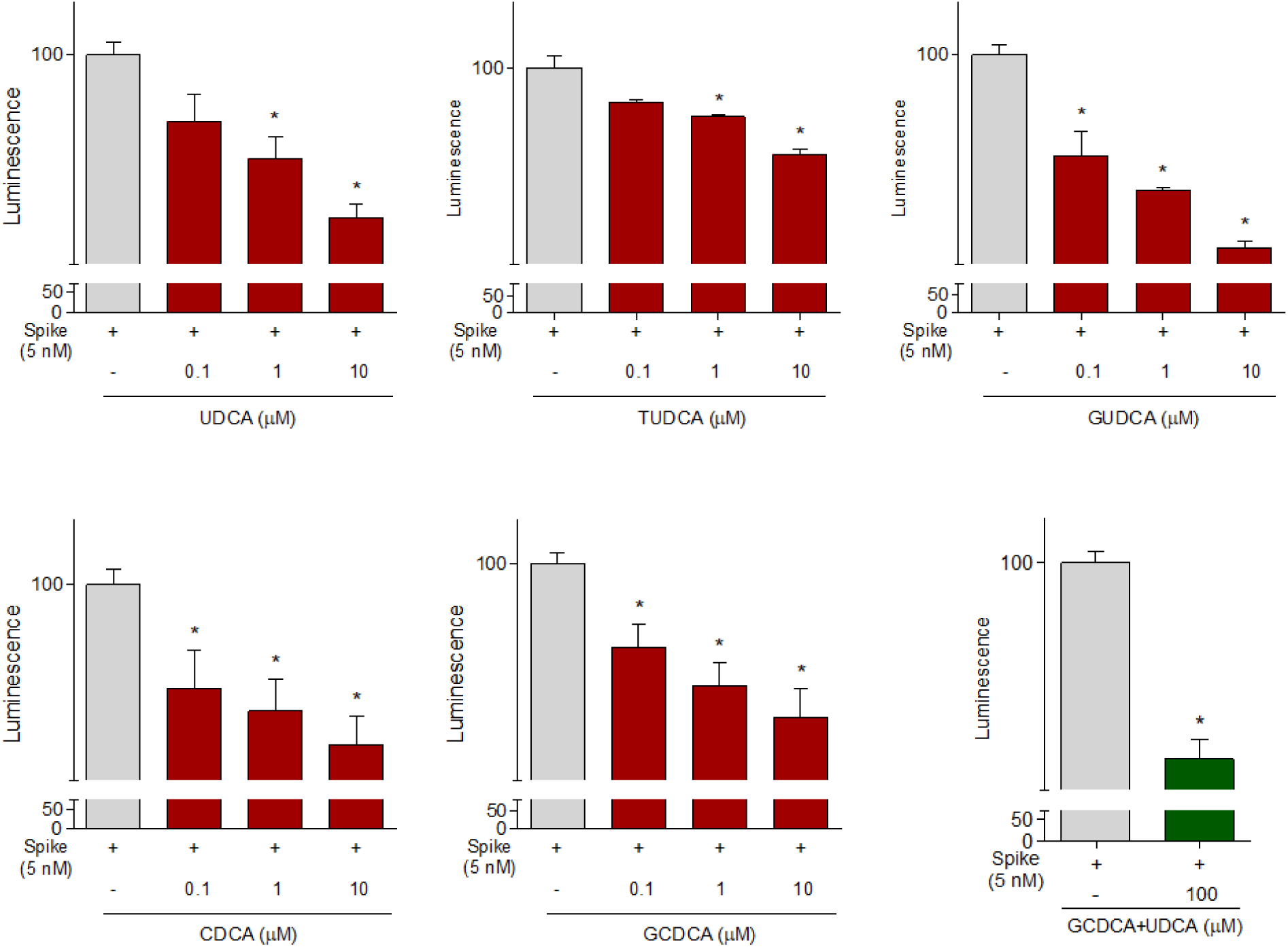
The ACE2:SARS-CoV-2 Spike Inhibitor Screening assay was performed as described in Material ad Method section. Natural bile acids UDCA, TUDCA, GUDCA, CDCA, GCDCA (0.1, 1 and 10 µM) and a combination of GCDCA + UDCA (100 µM), were tested to evaluate their ability to inhibit the binding of Spike protein (5 nM) to immobilized ACE2, by using the ACE2:SARS-CoV-2 Spike Inhibitor Screening assay Kit. Luminescence was measured using a Fluo-Star Omega fluorescent microplate reader. Luminescence values of Spike 5 nM were arbitrarily set to 100 %. Results are expressed as mean ± standard error. *p < 0.05 versus Spike 5 nM. Data are the mean ± SE, n = 3.

Continuing in the *in vitro* screening, we investigated whether the semisynthetic bile acid derivatives obeticholic acid (OCA), BAR704, BAR501 and BAR502, exert comparable or better effect than G-UDCA and G-GDCA. As illustrated in Figure 6, adding OCA to the incubation mixture reduced the binding of SARS-CoV-2 S spike to ACE2 by ≈20%. In contrast, BAR704, a 3-deoxy 6-ethyl derivative of CDCA, and a highly selective and potent FXR agonist was significantly more effective and reduced the binding by approximately 50% at the dose of 0.1 and 1µM. On the other hands, BAR501 and BAR502, alcoholic derivatives of UDCA and CDCA, respectively, were only slightly effective in reducing the binding of S protein RBD’s to ACE2.

**Figure 6.**
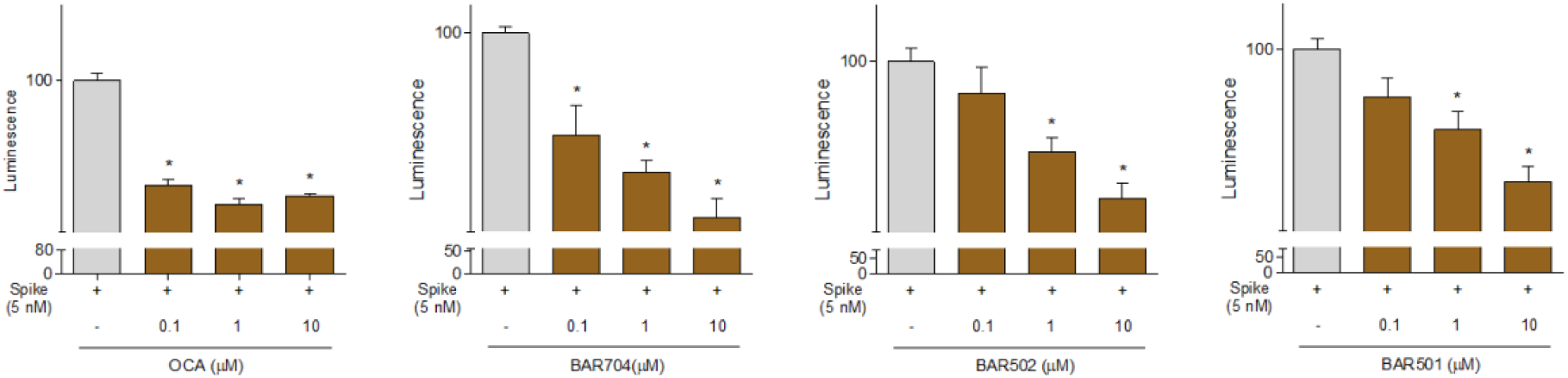
The ACE2:SARS-CoV-2 Spike Inhibitor Screening assay was performed as described in Materials and Methods section. The semi-synthetic bile acid receptor agonists OCA, BAR704, BAR502 and BAR501, were tested at different concentration (0.1, 1 and 10 µM) to evaluate their ability to inhibit the binding of Spike protein (5 nM) to immobilized ACE2, by using the ACE2:SARS-CoV-2 Spike Inhibitor Screening assay Kit. Luminescence was measured using a Fluo-Star Omega fluorescent microplate reader. Luminescence values of Spike 5 nM were arbitrarily set to 100 %. Results are expressed as mean ± standard error. *p < 0.05 versus Spike 5 nM. Data are the mean ± SE, n = 3.

### Effects of plasma samples from post-Covid-19 convalescent patients on Spike RBD –ACE2 interaction

To confirm the concept that binding the pockets in central β-sheet core of the Spike RBD effectively prevents its interaction with the consensus of ACE2 receptor, we have then carried out a set of control experiments using the plasma obtained from 5 patients that have recovered from Covid-19. These patients have a slightly different title of anti SARS-CoV2 antibody (See Material and Methods, Table 3), but all dilutions tested effectively inhibited the Spike RBD binding to ACE2 in our assay system by more than 95%. Together, these data highlight that the test used correctly identify the binding of SARS-CoV-2 RBD, but the level of inhibition, as expected was significantly lower than that could be reached by anti SARS-CoV2 antibodies.

**Table 3.** Percentage of inhibition of the Spike:ACE2 binding. Serum efficacy has been calculated in ACE2:SARS-CoV-2 Spike Inhibitor Screening Assay Kit as percent of inhibition of Spike RBD binding to ACE2 binding obtained using SPIKE at 5 nM, arbitrarily set as 100%.”

| Patient ID | Antibody Title | % of Binding Inhibition |  |  |
| --- | --- | --- | --- | --- |
|  |  | 5 μL of Serum | 10 μL of Serum | 20 μL of Serum |
| 1 | 96,6 AU/mL | 98,6 | 99,5 | 99,6 |
| 2 | 170 AU/mL | 99,3 | 99,4 | 99,3 |
| 3 | 89,4 AU/mL | 98,1 | 99,3 | 99,4 |
| 4 | 125 AU/mL | 98,8 | 99,3 | 99,4 |
| 5 | 146 AU/mL | 95,7 | 96,9 | 97,3 |

## Discussion

In this study we report the results of a virtual screening campaign designed to identify natural and clinically available compounds that might have utility in the prevention/treatment of SARS-CoV-2 infection. On the light of the need of effective therapies to be rapidly tested for preventing or treating the Covid-19, we have initiated an *in silico* campaign to identify putative molecular targets that could be exploited to prevent the interaction of the SARS-CoV-2 Spike protein with the cellular machinery hijacked by the virus to enter target cells. To this end, we have identified the Spike RBD as a potential pharmacological target. Accordingly, we have developed the concept that putative pockets on the surface of the central β-sheet core of the S protein RBD could be exploited eventually to prevent the binding of the virus to ACE2.

Our *in silico* screening has allowed the identification of six potentially druggable pockets and the virtual screening of the FDA-approved drug library identified steroidal compounds as potential hit against two pockets, namely pocket 1 and pocket 5. Interestingly, high accuracy docking demonstrated that flat steroidal scaffolds (i.e. AB junction *in trans* configuration Table 1) prefer pocket 1, while compounds with the A/B junction in *cis* configuration (Table 2, such as bile acids, show greater affinity for pocket 5.

Our *in vitro* testing has largely confirmed the functional relevance of the two main pockets identified by in silico analysis. One important finding of this study has been that several steroidal molecules were effective inhibitors of the binding of the RBD to ACE2 *in vitro*. In particular, the most interesting compounds in Table 1, the glycyrrhetinic and the oleanolic acid showed good agreements in terms of docking AD score and in their ability to inhibit the spike/ACE2 interaction *in vitro*. The results also suggested that the main determinant for the inhibition efficacy is the hydrophobicity, as demonstrated by the oleanolic acid, lacking any charge interaction within the pocket and resulting the most effective inhibitor in the series.

Hydrophobicity is also the main determinant of the activity of the bile acids and their semi-synthetic derivatives, as demonstrated by CDCA, the corresponding glyco-conjugated derivative (G-CDCA) and its semisynthetic derivatives OCA, BAR704, and BAR502. In fact, comparing the binding mode and the inhibition efficacy of CDCA and G-CDCA with the related 6-ethyl derivatives OCA and BAR704 highlighted the critical effect of the 6α-ethyl group in the inhibition activity. The above positive effect could be explained considering the internal energy contribution of these ligands to the ADscore, as well as the possibility of engaging more hydrophobic contacts. Indeed, the ADscore internal energy contribution, significantly higher for the 6-ethyl derivatives, represents a measure of the conformational energy of the bound *vs*. unbound state of the ligand, thus indicating that the ethyl group facilitates the assumption of the bioactive conformation. Moreover, the analysis of the binding mode of these compounds highlighted that the 6-ethyl in the α-position could establish hydrophobic contacts with P384 and Y369, positioned at a distance slightly longer than the optimal admitted for VdW interactions. However, it should be noted that the docking approach used considers the protein receptor as rigid and didn’t allow the mutual adaptation, which is an important process in ligand-receptor binding. In agreement with docking results, the lower efficacy observed for BAR502 could be explained with a slight change in the binding mode with a different position of the compound in the pocket in order to allow the hydroxyl group on a shortened side chain to interact with the side chain hydroxyl group of T376.

Moreover, also the comparison of the binding modes for G-CDCA and G-UDCA supported the hypothesis that the main determinant for the activity should be related to the network of hydrophobic interaction more than to the lack of a punctual hydrogen bond. Indeed, unlike the weakly active UDCA, the steroid core of G-UDCA is shifted to T376, and the resulting binding mode looks very similar to G-CDCA’s. Finally, the better inhibitory efficacy of BAR501 with respect to UDCA, further confirmed the not-essential effect of the charged group on the side chain in term of inhibition activity. Interestingly, the analysis of the binding mode of BAR501 also suggested that the stereochemistry of the ethyl group at C-6 is not pharmacophoric, being the 6β-ethyl group still able to potentially interact with P384 and Y369.

In the present study we have developed a strategy to target the interaction of SARS-CoV-2 S protein RBD with ACE2 receptor. As described in the introduction, SARS-CoV-2 enters the target cells by binging the carboxypeptidase domain of the ACE2 receptor, exposing a cleavage site, an hinge region between S1 and S2, to TMPRSSS2 which allows the S2 subunit of Spike protein to bind with cell membrane and virus penetration into host cells.

The two pockets we have identified in β-sheet core of the Spike RBD appears to be targetable by steroidal molecules and importantly we have found that either naturally occurring bile acids and their metabolites in human, i.e. G-UDCA and G-CDCA and T-UDCA and T-CDCA seem be able to reduce the binding of Spire RBD with ACE2. Among the natural and semi-synthetic agents, we have tested in this study, G-UDCA appears to be endowed with a robust activity toward Spike RBD and ACE2. Importantly we have found that most of the agents tested in this study were agonists to two main bile acid activated receptors, i.e. the Farnesoid-x-Receptor (FXR) and a cell membrane receptor known as GPBAR1. Thus, betulinic acid and oleanolic acid, along with UDCA, T-UDCA and G-UDCA, BAR501 and BAR502 are effective ligands for GPBAR1. In contrast, the glycyrrhetinic acid, CDCA, G-CDCA and T-CDCA, OCA and BAR704 are known for their ability to bind to FXR (Festa et al., 2014). The fact that all these agents bind to these two receptors and simultaneously to a viral protein was never rationalized before and might be exploited to reduce the hijack the potential of the virus to bind to the ACE2 receptor.

Of interest, some of these agents have been reported for the potential use as anti-HIV agents (Řezanka et al., 2009), while oleanolic acid that might function as a broad spectrum entry inhibitor of influenza viruses (Yang et al., 2018). On the other side, betulinic acid has been demonstrated useful in reducing inflammation and pulmonary edema induced by influenza virus (Hong et al., 2015), and potassium canrenoate, the main metabolite of spironolactone *in vivo*, is an anti-aldosteronic/diuretic used in the treatment of hypertension patients. Finally, several GPBAR1 and FXR ligands, as bile acid derivatives, have been proved to exert beneficial effect in immune disorders (Fiorucci et al., 2018) and among these, BAR501, the first example of C-6β-substituted UDCA derivative with potent and selective GPBAR1 activity, has been recently demonstrated a promising lead in attenuating inflammation and immune dysfunction by shifting the polarization of colonic macrophages from the inflammatory phenotype M1 to the anti-inflammatory phenotype M2, increasing the expression of IL-10 gene transcription in the intestine and enhanced secretion of IL-10 by macrophages (Biagioli et al., 2017).

In addition, we have found that other steroids, such as the potassium canrenoate has some ability to bind to the pockets of the Spike RBD thus limiting the binding of SARS-CoV-2 spike to ACE2 receptor.

One important observation we have made is that, while two different pockets of Spike RBD are potentially druggable, these are contiguous, and indeed when we have attempted several drug combinations, none of these combinations effectively increased the anti-adhesive efficacy of single agents.

This study has several limitations. First of all, we have observed that anti-adhesive effects exerted by hyperimmune plasma obtained from patients who have recovered from COVID-19, who have high titles of neutralizing antibodies, inhibit the Spike RBD/ACE2 interaction to a percentage close to 99%. This percentage is significantly higher than what we have measured with G-UDCA, the agent that gives the better results in this study. This suggests that small molecules binding of the hydrophobic pockets are less effective than a neutralizing antibody. This suggest that our pharmacological approach will likely be poorly effective in the presence of a highly replicative viral load. This also suggest that this approach might have some efficacy in preventing virus entry only in the case of low viral load. Nevertheless, the mild inhibition efficacy showed by c bile acids and their derivatives could pave the way for a further optimization of the binding mode exploring the chemical space of the steroidal nucleus in order to envisage additional potential interactions in pocket 5, recently demonstrated the least exposed to mutations in different circulating viral lineages.

Another limitation is that we have not tested the effect of these treatment on viral replication and further studies are needed to clarify this point.

In conclusion, in this paper, we have developed a strategy to target the interaction of SARS-CoV-2 S protein RBD with ACE2 receptor in identifying several steroidal derivatives as SARS-CoV-2 entry inhibitors. Our results demonstrate that several potential binding sites exist in the SARS CoV-2 S protein, and that the occupancy of these pockets reduces the ability of the S protein RBD to bind to the ACE2 consensus *in vitro*. Among all compounds screened in this study, natural occurring and clinically available steroids as glycyrrhetinic and oleanolic acids and bile acids derivatives have been proved to be effective in preventing virus entry in the case of low viral load. All together, these results might help to define novel approaches to reduce the viral load by using SARS-CoV-2 entry inhibitors.

